# Retrosplenial cortex and its role in spatial cognition

**DOI:** 10.1101/190801

**Authors:** Anna Mitchell, Rafal Czajkowksi, Ningyu Zhang, Kate Jeffery, Andrew Nelson

## Abstract

Retrosplenial cortex (RSC) is a region within the posterior neocortical system, heavily interconnected with an array of brain networks, both cortical and subcortical, that is engaged by a myriad of cognitive tasks. Although there is no consensus as to its precise function, evidence from both human and animal studies clearly points to a role in spatial cognition. However, the spatial processing impairments that follow RSC damage are not straightforward to characterise, leading to difficulties in defining the exact nature of its role. In the present article we review this literature and classify the types of ideas that have been put forward into three broad, somewhat overlapping classes: (i) Learning of landmark location, stability and permanence; (ii) Integration between spatial reference frames, and (iii) Consolidation and retrieval of spatial knowledge (“schemas”). We evaluate these models and suggest ways to test them, before briefly discussing whether the spatial function may be a subset of a more general function in episodic memory.

## Introduction

Retrosplenial cortex (RSC) has fallen within the scope of memory research for at least 40 years (Vogt, 1976) and yet as Vann et al. (Vann et al., 2009) pointed out in their recent comprehensive review, little was discovered about the structure for the first 90 years after Brodmann first identified it. Since the early 1990s, a growing body of evidence has implicated the RSC variously in spatial memory, navigation, landmark processing and the sense of direction, visuo-spatial imagery and past/future thinking, and episodic memory. Early results were difficult to interpret in the absence of precise neuroanatomical, behavioural, electrophysiological and functional data but as a consequence of intense research on the RSC, both across animal models using a variety of methods, and also in human neuropsychological and imaging studies, a group of theories is now emerging that highlight the involvement of the RSC in aspects of cognition that go beyond, yet at the same time still underlie, our abilities to process spatial information and retrieve memories. This review will examine the experimental data in light of its contribution to spatial cognition, beginning with a review of the anatomy and connectivity, followed by functional investigations based on lesion studies, imaging and electrophysiology, and concluding with evaluation and classification of the main ideas that have emerged. We suggest that the proposals about RSC function fall into at least three classes: first, that it is involved in the setting of perceived landmarks into a spatial reference frame for use in orientation (spatial and directional) as well as evaluation of landmark stability; second, that it stores and reactivates associations between different processing modes or reference frames for spatial navigation, and third, that it has a time-limited role in the storage and possibly retrieval of hippocampal-dependent spatial/episodic memories. We conclude with some suggestions about how to further refine, and perhaps ultimately synthesize, these models.

### Anatomy and connectivity of RSC

In human and non-human primates, RSC conforms to the cortical regions identified by Brodmann as areas 29 and 30, which - along with areas 23 and 31 - form part of the posterior cingulate cortex, lying immediately posterior to the corpus callosum (Fig. 1 A and B). Rodents lack areas 23 and 31, and RSC itself is located more dorsally and reaches the brain surface (Fig. 1C). Its central location makes it pivotally positioned to receive information from, and readily influence, many key brain regions responsible for the processing of spatial information.

**Figure 1.**
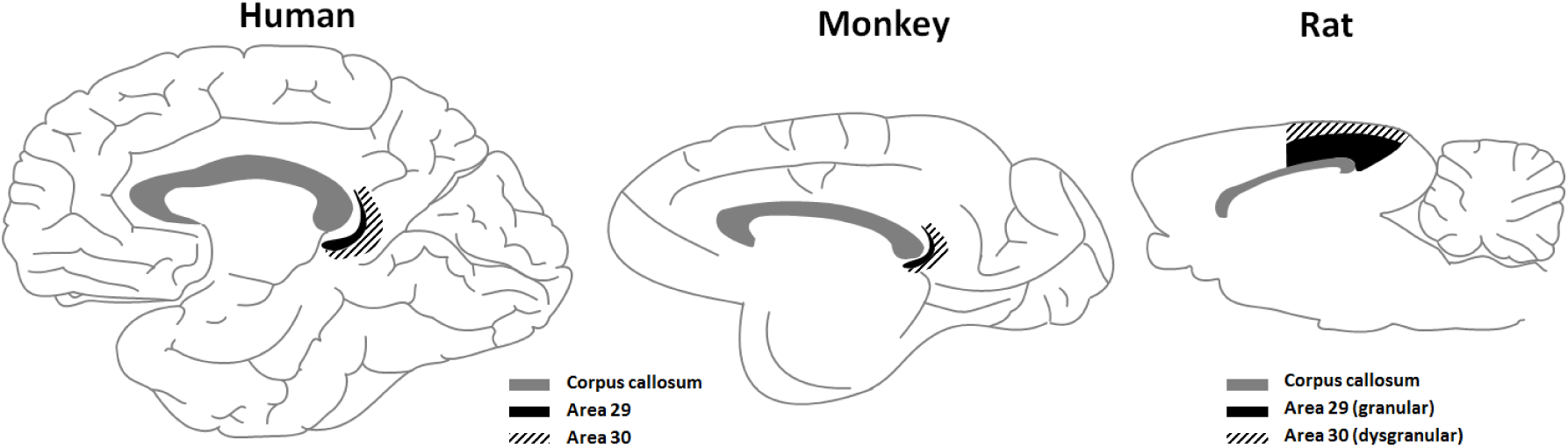
Schematic of the RSC as seen in midsagittal section and located just posterior to the corpus callosum, in humans, rhesus monkeys and rats. Figure by Jeffery, 2017; available at https://doi.org/10.6084/m9.figshare.5414179.v1 under a CC-BY 4.0 license.

Typically, structural neural connections have been mainly derived from studies in animal models (rodents and nonhuman primates), while the majority of neural connections studied in humans have been derived functionally. It is known that in both rats and primates, the majority of RSC (RSC granular A, and granular B, and RSC dysgranular) connections (up to 78%) originate in or are received from other parts of RSC, and from the posterior cingulate cortex in primates (Kobayashi and Amaral, 2003).

#### Cortical connections

As shown in Fig. 2, neural connections of the RSC from the cortex include the parahippocampal region (postrhinal cortex in rodents) (Suzuki and Amaral, 1994), medial entorhinal cortex (Van Hoesen and Pandya, 1975; Insausti et al., 1987; Jones and Witter, 2007; Insausti and Amaral, 2008; Czajkowski et al., 2013) and cingulate cortex (Jones et al., 2005). RSC receives unidirectional inputs from the CA1 field of the hippocampus (Cenquizca and Swanson, 2007; Miyashita and Rockland, 2007) and from the subiculum (Wyss and Van Groen, 1992; Honda and Ishizuka, 2015). It is also interconnected with extended hippocampal complex (including the presubiculum, postsubiculum and parasubiculum (Wyss and Van Groen, 1992; Kononenko and Witter, 2012), visuospatial cortical association areas (mainly medial precuneate gyrus, V4 of the occipital lobes, and the dorsal bank of the superior temporal sulcus) (Passarelli et al., 2017), and prefrontal cortex (with the heaviest terminations in the dorsolateral prefrontal cortex, frontopolar area 10, and area 11 of the orbitofrontal cortex); these frontal connections are all reciprocal. RSC also receives inputs directly from V2 of the occipital lobes. There are also prominent excitatory reciprocal connections between RSC and posterior secondary motor cortex - M2, recently identified in mice (Yamawaki et al., 2016).

**Figure 2:**
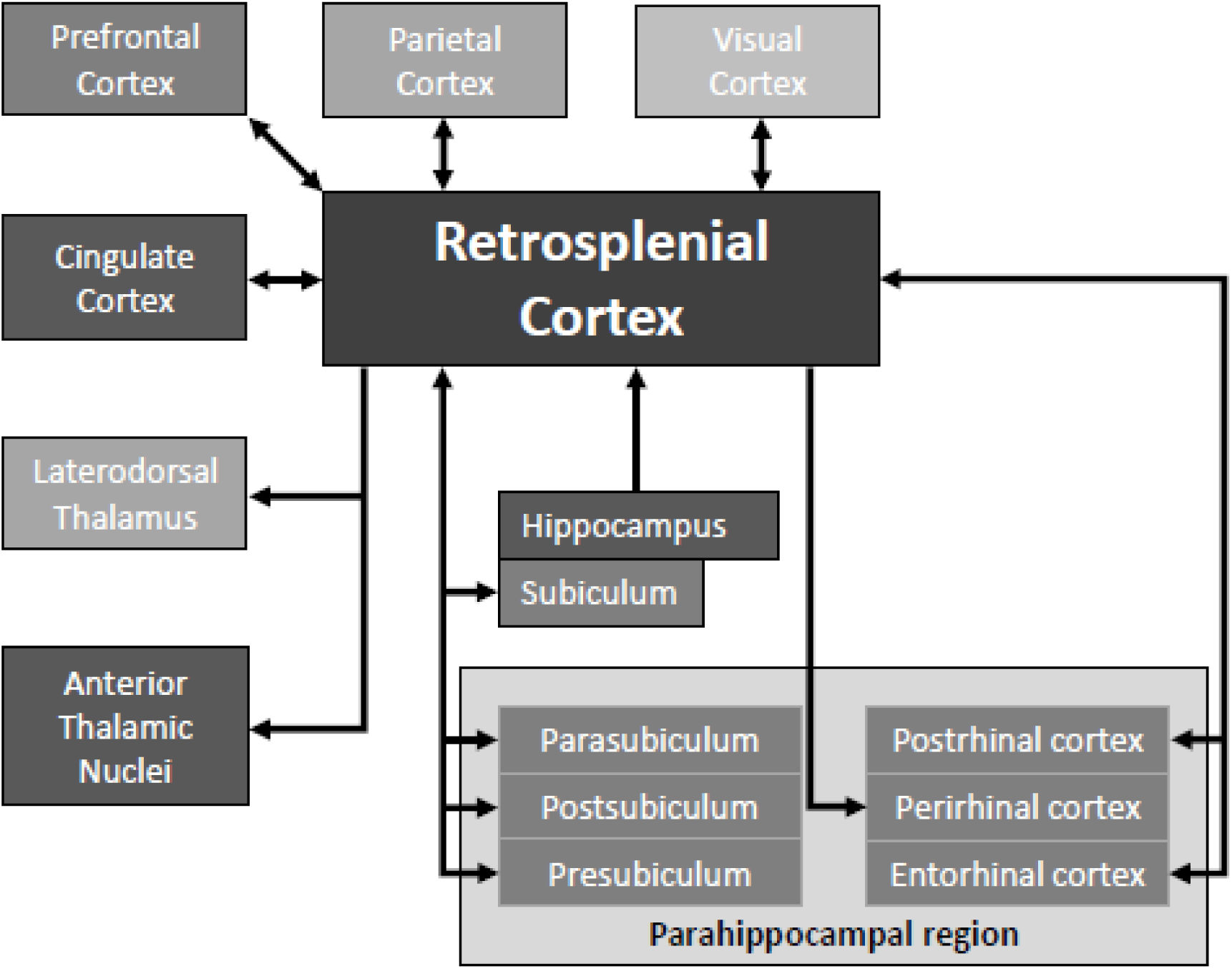
A schematic detailing the gross connectivity of retrosplenial cortex. As depicted in the figure, RSC serves as an interconnective hub for neocortical, hippocampal, parahippocampal and thalamic regions that are functionally involved in the processing of mammalian perceptions important for direction, location, landmarks and navigation.

#### Subcortical connections

In addition, as shown in Fig. 2, RSC has major reciprocal subcortical interactions with the anterior and laterodorsal thalamic nuclei (Vogt et al., 1987; van Groen and Wyss, 1990, 1992a, 1992b; Kobayashi and Amaral, 2003, 2007; Van Groen and Wyss, 2003; Aggleton et al., 2014) from where it receives heading direction information (Taube, 2007). There are also lesser connections with the mediodorsal thalamus and posterior thalamic nuclei (Powell, 1978; Aggleton et al., 2014) and additionally, RSC receives inputs from the intralaminar thalamic nuclei (important for arousal) and medial pulvinar (supporting visual attention) (Baleydier and Mauguiere, 1985; Vogt et al., 2006; Buckwalter et al., 2008).

In general, the anatomy shows that RSC interacts reciprocally with many brain regions, consistent with its role, described below, in a number of core cognitive competences. Of interest is the more unidirectional relationship with hippocampus, and with perirhinal cortex.

### Lesion studies

The literature on pure RSC lesions in humans is sparse and mostly from unilateral pathology, due to the rarity with which localized infarcts or injury occur to this region, and so most of our knowledge of human RSC comes from neuroimaging, which we discuss later. Most of the identified lesion-induced deficits appear to involve memory and spatial processing. Maguire (Maguire, 2001a) conducted a comprehensive review of the literature on RSC extant at the time and concluded that case studies of RSC lesions reveal deficits in episodic memory (memory for life events), occurring particularly following left-sided lesions, but also consistent reports of topographical disorientation (“getting lost”), with or without concomitant memory deficits, most of which followed right-sided lesions. The area that was most consistently involved in the pure disorientation cases was Brodmann’s area BA30. Maguire noted: “In every case, the patient was able to recognise the landmarks in their neighbourhoods and retained a sense of familiarity…Despite this, none of the patients was able to find his/her way in familiar environments, and all but one was unable to learn new routes”. Studies since then have confirmed the link between RSC lesions and topographic disorientation, with association of left-sided infarct with memory deficits (Kim et al., 2007), and of right-sided lesions with spatial impairment (Hashimoto et al., 2010, 2016), although spatial impairment has also been reported in patients with left-sided lesions (Ino et al., 2007; Ruggiero et al., 2014). Claessen and van der Ham conducted a review of lesion-related navigation deficits (Claessen and van der Ham, 2017) and found that involvement of the RSC was prominent in impairments of landmark-processing, and particularly in reporting distances and directions between known landmarks, or describing the positions of known landmarks or buildings on a map.

Experimental lesion studies in non-human primates can be much more precise, and also bilateral, which has provided new insights into RSC function. In rhesus macaque monkeys, damage to the RSC,which included the most caudal part of the posterior cingulate cortex, selectively impaired the ability to retrieve object-in-place scene discriminations that the monkeys had previously learnt (retrograde memory). In these tasks, animals have to learn and remember the location of a discrete object in a spatial scene. In contrast, these same animals were able to learn new object-in-place scene discriminations postoperatively (anterograde memory), so their ability to organise and retrieve spatial information remained intact. However, the 24-hour delay period between successive learning sessions of the new object-in-place discriminations (i.e. from session 1 to session 2) caused a specific impairment in retrieval of these new discriminations, resulting in the animals with RSC damage making more errors than controls during postoperative session 2 of new learning only. The task, object-in-place scene discriminations, incorporates elements of both spatial (e.g. landmark information) and episodic-like memory (unique object-in-place scene discriminations, with one of the objects in each discrimination paired with a reward if it is selected) without being explicitly autobiographical in nature (Gaffan, 1994; Mitchell and Gaffan, 2008; Mitchell et al., 2008; Murray and Wise, 2010). The novel findings observed in the monkeys’ performance led the authors to conclude that an intact RSC is particularly important for the ability to retrieve information that has been previously acquired, regardless of whether these memories are autobiographical, episodic (in the pure sense of what/ where and when) or actively spatial in nature (Buckley and Mitchell, 2016). Finally, the ability to retrieve this information did not require the monkeys to move around in their environment, although the successful execution of self-generated hand/arm movements (in order to select the correct object within the scene on the touchscreen) were necessary.

Studies involving rodents have proved vital in furthering our understanding of the contribution of the RSC to cognition, as they afford far greater neuroanatomical precision than is currently possible in primate studies. An early study by Berger et al. found that rabbits with RSC lesions could acquire a tone-light discrimination but were profoundly impaired in reversing it (Berger et al., 1986), suggesting a failure to modify a recently established memory. Given the dense interconnections between the RSC and the hippocampal spatial system, the majority of subsequent lesion studies have focused on spatial learning.

Some of the early studies into the effects of RSC lesions on spatial tasks produced mixed results. This divergence in findings may be attributable to methodological considerations such as the use of electrolytic or ablation lesions, which destroy fibres of passage and consequently may exaggerate the impact of the RSC damage; while other studies spared the more caudal aspect of the RSC, which is now known to be critically involved in spatial memory (Vann and Aggleton, 2002, 2004). Despite these earlier controversies, there is now very good evidence that RSC lesions in rodents disrupt spatial memory. Deficits are consistently reported on tasks that involve allocentric spatial processing, particularly when - as with the imaging studies - visual cues are needed for orientation (Hindley et al., 2014). Such tasks include learning the fixed or alternating location of a platform in the Morris water maze (Sutherland et al., 1988; Whishaw et al., 2001; Vann and Aggleton, 2002, 2004); the radial arm maze (Vann and Aggleton, 2004; Pothuizen et al., 2008; Keene and Bucci, 2009) and object-in-place discriminations (Parron and Save, 2004). There is some evidence that the RSC dysgranular region (area 30; see Fig. 1) may be particularly important for processing allocentric space, as rats with selective RSC dysgranular lesions were unable to use distal visual cues to guide spatial working memory, and relied instead on motor sequence information (Vann and Aggleton, 2005). Furthermore, deficits have also been found on tasks that require the use of directional information (Vann and Aggleton, 2004; Pothuizen et al., 2008; Keene and Bucci, 2009) as well as self-motion cues (Whishaw et al., 2001; Elduayen and Save, 2014). In some instances, the involvement of RSC has been found to be time-limited: for example, Maviel et al. (Maviel et al., 2004) found RSC inactivation in mice disrupted the retrieval of a recent one-day-old spatial memory but not a remote 30-day-old one, while Keene and Bucci found large impairments on radial maze performance for a 30s delay relative to a 5 s delay (Keene and Bucci, 2009). Findings such as these, combined with the IEG findings described later, suggest a particular role for RSC when spatial information needs to be retrieved from memory.

In general, the magnitude of spatial deficits after RSC lesions tends to be smaller and less striking than the spatial impairments associated with either hippocampal or anterior thalamic damage. Arguably, the classic demonstration of this difference is T-maze alternation performance, which is acutely sensitive to both hippocampal and anterior thalamic damage (Aggleton et al., 1986, 1996), but is often spared after RSC lesions (Meunier and Destrade, 1988; Neave et al., 1994; Pothuizen et al., 2008; Nelson et al., 2015b). Indeed, the full impact of RSC lesions often only emerges under specific conditions or when animals are required to shift between different spatial metrics. For example, temporary inactivation of the RSC selectively impairs navigation in the dark but not the light (Cooper et al., 2001). On the other hand, Wesierska et al. (Wesierska et al., 2009) found that rats with RSC dysgranular lesions could learn to avoid the shock zone of a rotating platform *if* the rotation occurred in the dark, so darkness per se does not seem to be the problem. The rats could also learn to avoid the shock zone if this was defined by allocentric room cues provided there were no conflicting local cues; thus there was not a straightforward impairment of allocentric cue use either. There *was* a notable impairment when the animals had to disregard the local cues and focus on the room cues. Thus, as the authors noted, impairments arose when relevant and irrelevant cues needed to be segregated. Similarly, impairments on both the radial arm maze and T-maze often only emerge when intra-maze cues are placed in conflict with extra-maze cues (Vann and Aggleton, 2004; Pothuizen et al., 2008; Nelson et al., 2015b).

A further illustration of the selective nature of RSC lesion-induced spatial deficits comes from an experiment by Nelson et al. in which the location of a submerged platform in a Morris watermaze was determined by either the geometric properties of the test environment or the juxtaposition of highly salient visual cues (Nelson et al., 2015a). Rats learnt the location of the platform by actively swimming to the platform or passively, that is by being repeatedly placed on the platform location prior to a test in which the animals had to swim to the correct location for the first time. RSC-lesioned rats were selectively impaired in the passive condition, indicating that RSC damage did not disrupt navigation per se but selectively impaired the ability to switch spatial frames of reference and different spatial viewpoints when navigating to the platform from a novel position in the environment (Nelson et al., 2015a). Similarly, complete RSC or selective RSC dysgranular lesions disrupted the ability to recognise the layout of a room from different viewpoints (Hindley et al., 2014).

Taken together, RSC effects appear to depend on the extent to which task performance relies on the use of spatial landmarks for orientation, or the need to switch between different spatial strategies or viewpoints. This is in line with the proposal that integration of the context in which an event occurs, learning about the significance of such stimuli or updating representations as new information comes on-line are all key aspects of RSC functioning.

### Brain imaging (PET, fMRI and IEG)

As outlined above, human, primate and rodent lesion studies have pointed to a role in spatial processing: complementary evidence comes from research using metabolic brain imaging, particularly positron emission tomography (PET), functional magnetic resonance imaging (fMRI) and immediate-early gene activation (IEG) studies. In an early PET study of cerebral glucose metabolism, Minoshima et al found a reduction in the posterior cingulate in patients with early Alzheimer’s disease, suggesting a role for this area in learning and memory (Minoshima et al., 1997). Nestor et al found that the RSC part of posterior cingulate was the most consistently hypometabolic region and suggested a role for this region in episodic memory retrieval (Nestor et al., 2003).

Since the advent of fMRI in cognitive neuroscience, many studies have investigated RSC activation as subjects perform tasks in the scanner. Indeed, RSC is now considered to be part of the so-called “default mode network”, which consists of a set of brain structures including medial frontal and medial temporal lobe regions, lateral and medial parietal areas and the RSC (Vann et al., 2009) that are active when subjects are not performing a task in the scanner, but rather are lying in the scanner at ‘rest’, or when they are actively simulating a situation (particularly one close in time and space to the present; (Tamir and Mitchell, 2011)) or when they are retrieving a memory (Sestieri et al., 2011).

Cognitive tasks that reliably activate RSC in fMRI studies include most that have a spatial component, especially when this requires use of the visual environment to retrieve previously learned information. These typically involve virtual reality simulations in which subjects navigate, by joystick or sometimes just by imagination, around a virtual environment, such as a town. In one of the earliest studies, Wolbers and Buchel scanned subjects as they learned a virtual maze-like town and found that RSC activation increased steadily with learning, and paralleled increasing map performance (Wolbers and Büchel, 2005). Similarly, in a study of London taxi drivers (Spiers and Maguire, 2006) in a virtual environment based on real maps of London, RSC activation occurred during route planning, spontaneous trajectory changes, and confirmation of expectations about the upcoming features of the outside environment (but not, interestingly, expectation violations). Another fMRI study confirmed that RSC activity was specifically associated with thoughts of location and orientation, as opposed to context familiarity or simple object recognition (Epstein et al., 2007). In both studies, the overall pattern of RSC activation differed from the one observed for hippocampus (Iaria et al., 2007), with the entire RSC active during both encoding and retrieval of spatial information.

A related line of work has investigated the encoding of location and/or direction by RSC. Marchette et al. (2014) performed multi-voxel pattern analysis (MVPA) of fMRI brain activation patterns on subjects recalling spatial views from a recently-learned virtual environment. Because MVPA compares fine-grained patterns of activation, it allows inferences to be made about whether a subject is discriminating stimuli. The virtual environment comprised a set of four museums located near each other in a virtual park. RSC activity patterns were similar when subjects faced in similar directions and/or occupied similar locations within each museum, suggesting that RSC was activating the same representations of local place and local direction, even though the environments were separated in global space. Similarity judgment reaction times were faster for homologous directions or locations, suggesting encoding by local features independent of global relationships. However, it was not demonstrated that subjects had been able to form global maps of the virtual space (that is, the reference frame in which the local spaces were set), so the question remains unanswered about whether RSC is also involved in encoding global space.

Robertson et al. (2016) also found encoding of local “landmarks” in a setting in which subjects viewed segments of a 360-degree panorama that either did or did not overlap. RSC activation was higher when subjects subsequently viewed isolated scenes from the overlap condition and judged whether it came from the left or the right side of the panorama. A study by Shine et al. (2016) did, however, find evidence for global heading representation in RSC. They investigated RSC and thalamus activation in subjects who had learned a virtual environment by walking around with a head-mounted display, which provides vestibular and motor cues to orientation, and found activation of both structures, which both contain directionally sensitive head direction cells (discussed below), when subjects were shown stationary views of the environment and had to make orientation judgments (Shine et al., 2016). Furthermore, recent work has also examined RSC activation when participants navigate in a virtual 3D environment (Kim et al., 2017). Interestingly in this study, the RSC activation was particularly sensitive to the vertical axis of space, which the authors suggest may be supporting processing of gravity, which is a directional cue in the vertical plane, and may be useful for 3D navigation. Given that there is evidence for both local and global encoding of direction in RSC, the question arises as to how these might both be accommodated within the one structure; we return to this question later.

While the foregoing studies looked at global spatial environments, work from the Maguire lab has suggested a role for RSC in the processing of individual landmarks. Auger et al. (2012) scanned subjects as they viewed a variety of images of large objects and found that RSC was activated only by the spatially fixed, landmark-like objects, and furthermore that the extent of activation correlated with navigation ability. In a follow-up study using multi-voxel pattern analysis, Auger and Maguire (2013) showed that decoding of the number of permanent landmarks in view was possible, and more so in better navigators, concluding that “RSC in particular is concerned with encoding every permanent landmark that is in view”. They then showed that this RSC permanence encoding also occurred when subjects learned about artificial, abstract landmarks in a featureless “Fog World” (Auger et al., 2015), demonstrating that the RSC is involved in new learning of landmarks and their spatial stability, and also that such learning correlates with navigation ability (Auger et al., 2017).

Some meta-analyses of human imaging studies have indicated that higher RSC activation occurs when subjects process landmark information (Maguire, 2001b; Spiers and Maguire, 2006; Auger et al., 2012; Mullally et al., 2012; Auger and Maguire, 2013) and associate the current panoramic visual scene with memory (Robertson et al., 2016). Further evidence has revealed that RSC is activated when subjects retrieve autobiographical memories (Maddock, 1999; Spreng et al., 2009) or engage in future thinking or imagining (Tamir and Mitchell, 2011), although RSC appears more engaged with past than future spatial/contextual thinking (Gilmore et al., 2017). While the retrieval of autobiographical memories, imagining and future thinking may not explicitly engage spatial processes, they are nonetheless closely allied to the spatial functions of RSC, as they require self-referencing to spatial contexts and the updating of spatial representations as events are recalled or imagined.

Animal models, in particular rodent experiments which engage their ability to readily explore their spatial environment, have provided more conclusive evidence highlighting the importance of the RSC for spatial functioning. One particular experimental approach is to study RSC functioning in the intact rodent brain by investigating the extent of and location of activation of learning-induced immediate early genes (IEG: e.g., *Arc, Fos* or *Zif268*) after animals have performed a behavioral task. Most of these studies have shown increased expression of IEGs as a consequence of spatial learning (Vann et al., 2000; Maviel et al., 2004), confirming the importance of plasticity within RSC for spatial memory. One of the distinct advantages of this approach is that it allows for far greater anatomical precision, for example revealing subregional or layer-specific difference in RSC activity after animals have performed a spatial task (Pothuizen et al., 2009).

IEG studies have also revealed the involvement of RSC involvement in spatial memory formation. Tse at al. (Tse et al., 2011) investigated the two IEGs *zif268* and *Arc* as rats learned flavour-place pairs; they found upregulation of these genes in RSC when animals added two new pairs to the set. A more advanced approach has been to combine IEG mapping with chronic *in vivo* two-photon imaging to study the dynamics of *Fos* fluorescent reporter (FosGFP) in RSC dysgranular during acquisition of the watermaze task (Czajkowski et al., 2014). Higher reporter activity was observed when animals relied on a set of distal visual cues (allocentric strategy), as compared to a simple swimming task with one local landmark. Moreover, these observations also revealed a small population of neurons that were persistently reactivated during subsequent sessions of the allocentric task. This study showed that plasticity occurs within RSC during spatial learning, and also suggested that this structure is critical for formation of the global representation. Indeed, in another set of experiments, optogenetic reactivation of *Fos*-expressing neuronal ensembles in mouse RSC led to replication of context-specific behavior in a safe context, devoid of any features of the original training context (Cowansage et al., 2014).

Taken together, these complementary human and animal studies have highlighted that RSC functioning is involved in spatial learning and memory, particularly when environmental cues (landmarks) are to be used for re-orientation and perhaps navigation. Studies of the time course of RSC involvement suggest a dissociation between new learning and memory retrieval/updating. The implication is that RSC is less involved in spatial perception per se, and more involved with memory retrieval and editing.

### Single neuron studies

Researchers typically turn to rodent single-neuron studies to address fine-grained questions about encoding. The first electrophysiological studies of spatial correlates of rodent RSC were conducted by Chen et al. (Chen et al., 1994a, 1994b) who reported that around 10% of RSC cells in the rat have the properties of head direction (HD) cells. These are cells that fire preferentially when the animal faces in a particular global direction; cells with these properties are found in a variety of brain regions, and are thought to subserve the sense of direction (Taube, 2007). However, 90% of the RSC neurons had more complex firing correlates, and no clear hypothesis about overall RSC function emerged.

A later study by Jacob and colleagues (Jacob et al., 2017) similarly found a subpopulation of HD cells, in the RSC dysgranular region only, the firing of which was controlled by the local environmental cues independently of the global HD signal, and – also like Chen et al. – a further sub-population of cells that showed mixed effects. This experiment took place in an environment composed of two local sub-compartments (two connected rectangles) that had opposite orientations within the room as a whole. Some cells behaved like typical HD cells and fired whenever the animal faced in a particular direction in the global space, while others fired in one direction in one compartment and the opposite direction in the other compartment ‒ as if these cells were more interested in local direction than global direction. This observation is thus reminiscent of the fMRI experiment by Marchette et al, discussed earlier, in which human subjects showed similar RSC activation patterns in local subspaces independent of their global orientation. Together, these results support the idea that RSC might be involved in relating spatial reference frames, with some cells responsible for local orientation and others responsible for the “bigger picture”.

More broadly, the findings concerning HD cells suggest that RSC neurons may be integrating landmark information, coming from the visual cortex, together with the ongoing head direction signal being assembled and maintained by more central elements, or nodes, in the HD network. Such interaction might depend on the strength and/or reliability of the sensory input (i.e. landmarks) to RSC and/or the HD system (Knight et al., 2014), raising the possibility that RSC directional neurons have the task of evaluating landmarks and deciding whether they are stable and/or reliable enough to help anchor the sense of direction (Jeffery et al., 2016).

The above notwithstanding, only around 10% of RSC neurons seem to be HD neurons, the remainder having more complex firing correlates. Many of these seem related to the actions the animal is performing. The first systematic analysis by Chen et al (Chen et al., 1994a, 1994b) reported RSC cells related to body turns in addition to those with spatial firing characteristics. A subsequent study found RSC cells with firing significantly correlated with running speed, location and angular head velocity (Cho and Sharp, 2001). Similarly, cells that respond to specific combinations of location, direction and movement were reported by Alexander and Nitz (Alexander and Nitz, 2015) who recorded RSC neurons as rats ran on two identical ‘W’-shaped tracks located at different places in a room. As well as ordinary HD cells (here around 6%), they found cells encoding conjunctions of local position, global position and left/right turning behaviour. In a later study (Alexander and Nitz, 2017), some RSC neurons were found to show firing rate peaks that recurred periodically as animals ran around the edge of a plus maze ‒ some cells activated once per circumnavigation, some twice, some four times etc. This observation again suggests a possible role in relating local and global spatial reference frames. However, recurring activation patterns were also seen when the animal ran on a ring track, with no local substructure, so it is possible that the cells were responding to some type of fourfold symmetry in the distal room cues.

In contrast to encoding of route, within which every location that the animal visits along the full trajectory is represented, others have reported encoding of navigational or behaviourally significant cues (e.g. goal-location coding) by RSC in simpler linear environments. In a study by Smith et al. (Smith et al., 2012), animals on a plus-maze learned to approach the east arm for reward for half of each session and then switched to the west. RSC neurons developed spatially localised activity patches (‘place fields’) that were sensitive to reward-associated locations, and the number of place fields substantially increased with experience. However, unlike co-recorded hippocampal place cells, which produce very focal place fields, RSC place fields were dispersed and sometimes covered the entire arms. One function of RSC place fields could be enabling the rats to discriminate two behavioural contexts.

A recent study by Mao et al. (Mao et al., 2017) reported more hippocampal-like activity in RSC cells, finding spatially localised activity (i.e., place fields) on a treadmill during movements in head-fixed mice. Locations on the track were marked by tactile cues on the travelling belt. As with hippocampal place cells, changes in light and reward location cause the cells to alter their firing locations (“remap”). These observations support the notion that RSC is sensitive to spatially informative cues and contextual changes.

In addition to place, cue-, and reward-location, conjunctive coding was reported by Vedder et al. (Vedder et al., 2016). In a light-cued T-maze task, RSC neurons increased responsiveness to the light cue, mostly irrespective of left-right position, but they also frequently responded to location or to reward. Responses involved both increased firing (“on” responding) and decreased firing (“off’ responding). Interestingly, responding to the light often slightly preceded the onset of the light cue, an anticipatory response also reported earlier in RSC in rabbits in an associative conditioning task (Smith et al., 2002). The location-sensitive firing on the stem of the T often distinguished forthcoming left and right turns, so-called “splitter” behaviour also seen in hippocampal place cells (Wood et al., 2000). Thus, although responding is associated to a cue, there seems sometimes to be a supra-sensory component related to expectation.

To summarize, then, the results from electrophysiology studies of RSC neurons provide a mixed picture in which spatial processing dominates, but the nature of the processing is hard to pin down exactly. It is clear that place and heading are represented, but so are other variables, and the nature and function of the conjunctive encoding remains unclear.

### The RSC contribution to spatial cognition - consensus and controversies

The experimental literature reviewed above has revealed areas where investigators are in general agreement, and other areas where there is debate or uncertainty. In this section we review these areas and outline some ways forward to resolve these.

There seems to be general agreement that RSC has a role in allocentric spatial processing; where there are differences is in the exact contribution it makes, and also in whether it has a broader role in memory, of which space is just a subcomponent. In this regard, the status of RSC research is a little reminiscent of hippocampal research 30 years ago. Our conclusion is that the literature has yielded three broad, somewhat related views concerning RSC’s spatial function:

(i) That it processes landmarks and landmark stability/permanence, in service of spatial/directional orientation
(ii) That it mediates between spatial representations, processing modes or “reference frames”
(iii) That it is involved in consolidation and retrieval of spatial schemas, for example to support episodic memory

#### Landmark processing

The first set of views is that RSC has a specific function in the encoding of the spatial and directional characteristics of landmarks, independent of their identity. This view emerges from such findings as that HD-cell sensitivity to landmarks is reduced following RSC lesions (Clark et al., 2010), that some RSC directionally tuned cells respond to environmental landmarks in preference to the main HD network signal (Jacob et al., 2017), that RSC is active when humans process landmark permanence (Auger et al., 2012, 2017; Auger and Maguire, 2013) and that lesions to RSC in human subjects cause them to lose the ability to use landmarks to orient (Iaria et al., 2007). It is also supported by findings that rats with RSC lesions are poor at using allocentric spatial cues to navigate (Vann and Aggleton, 2005). By this view, the function of RSC is to process landmarks as currently perceived, and fit them to a spatial framework so that in future they can be used for self-localisation and re-orientation. This viewpoint supposes a particular role for landmarks in the formation of spatial representations, and is consistent with the close relationship of RSC to visual areas as well as to the hippocampal spatial system.

#### Spatial representations

The second view, which could be regarded as an extension of the first to information beyond landmarks, is that this area serves to mediate between spatial representations. For some investigators, this has meant between egocentric and allocentric processing, although “egocentric” suggests different things to different researchers, meaning self-motion-updated to some, and viewpoint-dependent to others. Chen et al. (1994a) made the specific proposal that the egocentric information concerns self-motion, a position also taken by Alexander and Nitz (Alexander and Nitz, 2015); this is supported by their and others’ observations that directionally tuned neurons in RSC are updated by self-motion (sometimes called idiothetic) cues and that navigation in RSC-lesioned animals is affected by darkness (Cooper and Mizumori, 1999). Other authors have taken “egocentric” more broadly to mean spatial items encoded with respect to the body vs. with respect to the world. For example, Burgess and colleagues have suggested that RSC is part of progressive cortical transform of parietal egocentrically to hippocampal allocentrically encoded information (Byrne et al., 2007), and the hypothesis of egocentric-allocentric transformation by RSC recurs repeatedly in the literature (see Vann et al 2009 for discussion).

Another, not dissimilar view is that RSC is involved in constructing and relating allocentric spatial reference frames more generally, not necessarily egocentric/allocentric ones. This view is supported by studies such as the museums study of Marchette et al. (Marchette et al., 2014) which found similarities in the encoding of local spaces even though these were separated in global space, and the related findings of Jacob et al. (Jacob et al., 2017) that some RSC neurons constructed a directional signal based on local cues while others used the global space. A related notion is that RSC is involved in switching between different modes (as opposed to frames) of spatial processing - from light to dark (Cooper and Mizumori, 1999; Cooper et al., 2001), or distal to proximal cues etc.

These models all share the underlying feature that RSC has access to the same spatial information represented in different ways, and is needed in order to switch between these.

#### Spatial schema consolidation/retrieval

The final view, which is broadest of all, is that RSC is involved in formation and consolidation of hippocampus-dependent spatial/episodic memories. What differentiates these models from the foregoing, and also from standard theories of hippocampal function, is the incorporation of a temporal dimension to the encoding. By this is meant that RSC is not needed for de novo spatial learning, but *is* required when the animal needs to draw on a previously learned set of spatial relationships, in order to execute a task or acquire new information to add to its stored representation.

These ideas draw on two sets of theoretical work already extant in the literature: the idea proposed by Marr (Marr, 1971) that rapidly formed hippocampal memories are slowly consolidated in neocortex, and the idea that spatial learning entails the formation not just of task-or item-specific memories, but also of a more general framework within which the memories are situated, which has sometimes been called a schema (Ghosh and Gilboa, 2014). An example of a schema might be the watermaze task, in which a rat is faced with needing to learn a new platform location: learning of the new location is faster than the original because the rat already knows the layout of the room and the watermaze, and the procedures required to learn the platform location - learning the location for *today* requires just a small updating.

Support for this consolidation/updating idea comes from multiple observations in the literature that the role of RSC in behavioural tasks is frequently time-limited in that its effects occur later in training rather than immediately. In particular, there seems to be a 24-hour time window after training, below which RSC is engaged less, but after which it becomes involved: see the experiment by (Buckley and Mitchell, 2016), and the observation by Bontempi and colleagues that IEGs are upregulated when mice retrieve a 30-day-old memory but not a one-day-old one (Maviel et al., 2004). IEG studies have also revealed greater engagement for RSC in spaced vs. massed training (Nonaka et al., 2017), consistent with the need, in spaced training, for reactivation of a partly consolidated (10 min-old) memory rather than a completely newly formed (30 s-old) one.

This time-dependency has led several investigators to propose that RSC is part of a “primacy system” (Gabriel, 1990; Bussey et al., 1996) the function of which is to retrieve and process information learned earlier. One of the most detailed of such models was put forward by Ranganath and Ritchey (Ranganath and Ritchey, 2012) who proposed that RSC, together with parahippocampal cortex, forms part of a posterior cortical network that functions to support episodic memory. They suggest that this network matches incoming cues about the current behavioural context to what they call “situation models” which are internally stored representations of the relationships among the entities and the environment. According to their view, the parahippocampal cortex identifies contexts and the RSC compares these external cues with internal models of the situation, including input regarding self-motion.

### Open questions

Resolving the above ideas into a single, inclusive model of RSC function (if this is possible) will require the answering of some of the outstanding questions raised by the studies to date. Below, we outline some of these questions.

1. Does RSC have a specific interest in landmarks, as a subclass of spatial cue? A finding that has emerged from multiple studies of spatial processing is that RSC is particularly involved in the processing of landmarks, which is to say discrete objects or visual discontinuities in the panorama that serve, by virtue of their distant location and spatial stability, to orient the sense of direction. However, the specific hypothesis that it is interested in landmarks as discrete objects as opposed to, say, visual panoramas, has not been fully tested. An unanswered question then is whether RSC is engaged during spatial processing in the *absence* of landmarks; for example in an environment devoid of discrete spatial cues in which geometry or smooth visual shading provide the only cues to direction. It should be noted that most types of geometric environment (squares, rectangles, teardrops etc) have corners, which could in principle act as discrete landmarks, so care would have to be taken with the environmental design to ensure the absence of all such discrete visual stimuli. The general question to be answered here is whether the brain, via the RSC, treats landmarks as a special category of object or whether the interest of RSC in landmarks stems solely from their spatial utility, which in turn derives from the constant spatial relations between them.
2. Does RSC mediate between spatial representations? The core idea here is that RSC may not be needed for spatial learning per se, but *is* needed when the subject moves between representational modes. This may entail switching from egocentric to allocentric encoding of cues, or relating an interior space to an exterior one (for example, deciding which door one needs to exit through to reach the carpark). This view is an extension of the local idea discussed above, that RSC is needed to be able to use spatial landmarks to retrieve current location and heading. The important new ingredient supplied by the reference frame framework, as it were, is that at least two representations have had to be activated: for example, being in one place and thinking about another, or navigating in the dark and remembering where things are based on experience in the light. The question to be answered, therefore, is whether RSC is indeed needed for a subject to activate two representations simultaneously. Testing this idea is complicated by the demonstration discussed earlier that RSC is also needed to use local spatial cues to retrieve a previously learned spatial layout. Since it is required for current self-localisation, the idea that it is also needed for spatial imagination, or route planning, or future thinking, or other similar imagination-based functions, is hard to test directly because a lesioned subject can’t even get past the initial orientation problem. However, more temporally focused interventions such as optogenetic (in)activation may be of help here. For example, once intact animals have learned a radial maze task, it may be found that RSC is needed if the lights are turned off halfway through a trial, forcing a switch from one processing mode to another. Similarly, rats familiarized with a small space inside a larger one may be able to navigate between the two when RSC is operating, but not when it is inactivated. These types of task probe retrieval and manipulation of already-stored spatial representations.
3. What is the time course of RSC involvement in spatial learning? It remains an open question exactly when RSC comes into play during formation and use of spatial memories. IEG studies have found that activation occurs rapidly (within minutes) of modifying a spatial schema (Tse et al., 2011), and the massed/spaced learning experiment from the same group suggests a role for RSC during at least the first few minutes (Nonaka et al., 2017). While other animal studies have found that RSC dependence of a task does not appear until memories are reactivated again after a delay of up to 24 hours (e.g., (Buckley and Mitchell, 2016), presumably to allow an epoch of time to pass before re-engagement of the RSC memory can occur. Consequently as our current understanding stands, it is difficult to distinguish if there is a critical time window in which RSC is engaged by spatial tasks or whether methodological issues, such as divergences in experimental design or even species specific differences, can explain the discrepancies in the literature. Nevertheless, this issue can readily be addressed empirically. One approach would be to take a task that is known to induce the expression of IEG within RSC and compare the effects of blocking IEG expression with antisense oligonucleotides at different stages of task acquisition (early versus late stage). Pharmacological or chemogenetic silencing of RSC neuronal activity could similarly be used to assess whether the RSC is differentially involved in remote or recent spatial memory (Corcoran et al., 2011). Studies in rodents can be complemented by imaging studies in humans that compare RSC activity in participants navigating in new or previously learnt virtual or real environments (Patai et al., 2017).
4. What is the relationship between RSC and hippocampus? RSC first attracted attention because of its links with the hippocampal memory system, and as discussed here, many of the deficits arising from RSC damage resemble those of hippocampal lesions, with some notable differences. Of interest is the asymmetric relationship with hippocampus, in that RSC receives more direct connections (from CA1 and subiculum) than it sends, although it does project indirectly to hippocampus via entorhinal cortex and the subicular complex. It will thus be important to determine the interaction between these structures, during memory formation, retrieval and updating. Targeted optogenetic interventions will be useful here, as they allow selective interruption of pathways (e.g., those afferents that came from hippocampus alone), and temporally precise interruption of processing epochs. The extent to which RSC and hippocampus are functionally coupled could also be examined by combining temporary inactivation techniques with electrophysiology, for example assessing the effect of hippocampal inactivation on neuronal firing within RSC or vice versa. Extant data suggests that RSC inactivation causes changes in hippocampal place fields (Cooper and Mizumori, 2001) but many questions remain open. By combining the latest chemogenetic techniques with electrophysiological recordings and behavioural assays, researchers will be able to address questions about the nature of functional interactions between RSC and hippocampus with far greater anatomical and temporal precision. Future studies could also explore whether RSC and hippocampus are engaged by different navigational strategies, such as map-based, route-planning, and scene construction. Furthermore, RSC may work separately from the hippocampus in processing previously consolidated spatial information (see above), as evident in a recent work by Patai et al (Patai et al., 2017). This study reported higher RSC activity during distance coding in familiar environments, in contrast to higher hippocampal activity seen in newly learned environments, where more route-planning might have occurred.
5. What is the role of RSC in episodic memory more broadly? A review of the human literature reveals a difference between left and right RSC in both lesion findings and imaging; in particular, the left seems to be more implicated in general episodic memory, while the right is more implicated in spatial processing. Is RSC also involved in episodic memory in animals? We still lack a good animal model of this form of memory, because most animal tasks require training whereas episodes are, by their nature, transient. Nevertheless it will be important to determine, in future, the extent to which RSC has a role in memory that extends beyond space. Indeed, evidence is now emerging implicating RSC in mnemonic processes that do not contain any obvious spatial component including the processing or retrieval of temporal information (Todd et al., 2015; Powell et al., 2017) as well as learning the inter-relationship between sensory stimuli in the environment (Robinson et al., 2014); processes that are likely to be central to our ability to remember an event. Reconciling these seemingly disparate spatial and non-spatial roles, therefore, represents a key challenge for understanding RSC function.

### Summary and conclusions

In conclusion, we have reviewed the literature on the RSC contribution to spatial memory, and have found that there are three broad classes of models, which differ in their focus but have significant overlaps. It remains unclear whether RSC has more than one function, or whether these three classes of function are better described by some overarching model that can explain the current findings. We have outlined some open questions, the answers to which will require an interaction between multiple different approaches, in a variety of species.

Over one hundred years have passed since Brodmann first identified the RSC, and, while in the intervening years significant advances have been made in elucidating the role RSC plays in cognition, the precise functions of the RSC still remain somewhat of an enigma. It is hoped that the framework set out in this review will provide a basis for subsequent endeavours to probe the underlying function(s) of this most fascinating of brain structures.

## Acknowledgments

This work was supported by a Wellcome Trust Senior Research Fellowship in Basic Biomedical Sciences to A.S.M. (110157/Z/15/Z), a grant from National Science Centre (Poland) Sonata Bis 2014/14/E/NZ4/00172 to R.C., a scholarship from Chinese Scholarship Council to N.Z. (201608000007), a Wellcome Trust Investigator Award to K.J. (WT103896AIA) and grants from the BBSRC (BB/H020187/1 and BB/L021005/1) to A.J.D.N.

